# Novel 3D Mature Osteocyte Perfusion Platform Reveals Subtype-Dependent and Radiation-Sensitive Disruption of Mechanotransduction by Breast Cancer Secretomes

**DOI:** 10.64898/2026.10.22.760275

**Authors:** Antonio Salcido-Alcántar, Corrin Bowman, Ellie Goldman, Emma N Briggs, Anthony M Bierschenk, Tanvi K Rami, William R Thompson, Maureen E Lynch

**Author notes:** These authors contributed equally (Antonio Salcido-Alcántar, Corrin Bowman).

## Abstract

Osteocytes (OCys) coordinate bone remodeling by transducing mechanical signals into anabolic cues, primarily through suppressing sclerostin and RANKL. In breast cancer (BCa) bone metastasis, mechanical loading fails to restore lost bone, pointing to tumor-induced OCy mechanosensory dysfunction. Here, we investigate how BCa molecular subtype heterogeneity and frontline therapies like radiation influence OCy mechanobiology. We leveraged a 3D mineralized scaffold paired with a forced perfusion bioreactor to establish a physiologically relevant model of mature, dendrite-bearing OCys. After optimizing perfusion parameters, 3D mature OCys were exposed to conditioned media (CM) from estrogen receptor-positive (ER+; ZR-75-1) or triple-negative (TN; MDA-MB-231) BCa cells generated 24 h post-irradiation (0 vs. 2 Gy). Downstream OCy gene expression (*Sost, Tnfsf11, Tnfrsf11b*) and tumor CM proteomic profiles (84 oncology targets) were evaluated. Differentiated 3D MPC-2s exhibited interconnected dendritic networks, classical responsiveness to parathyroid hormone, and robust flow-induced *Sost* suppression. Non-irradiated ER+ BCa CM preserved canonical OCy mechanoresponsiveness (*Sost* downregulation post-flow), but this response was eliminated when ER+ cells received 2 Gy radiation. Conversely, TN CM blunted flow-induced *Sost* suppression regardless of irradiation status. Proteomic profiling revealed that BCa subtype was the primary driver of secretome composition, whereas 2 Gy radiation induced distinct directional secretome shifts between subtypes. These findings demonstrate that BCa-induced impairment of OCy mechanosensing is subtype-dependent and that low-dose radiation further exacerbates secretome-mediated mechanosensory loss in ER+ disease, highlighting key tumor-derived targets for restoring skeletal health.

## INTRODUCTION

Osteocytes (OCys) comprise over 90% of all bone cells and function as the primary mechanosensors and master orchestrators of skeletal homeostasis [1, 2]. They sense and interpret fluid shear stresses generated by physical loading to coordinate downstream bone remodeling. In healthy bone, skeletal loading stimulates bone anabolism primarily via OCy mechanotransduction, which upregulates osteoprotective/anabolic factors (e.g., OPG) and suppresses anti-anabolic signals such as sclerostin and RANKL. Mechanical loading is traditionally considered osteoprotective and an attractive non-pharmacological strategy to mitigate skeletal degradation, including loss caused by diseases such as bone metastasis. However, in bone-metastatic conditions such as breast cancer (BCa), in vivo loading (e.g., tibial compression [3-8], vibration [9], and treadmill running [7]) merely slows or mitigates bone destruction rather than fully preventing or restoring tumor-induced bone loss [4, 6, 9, 10]. OCy dysfunction is implicated in bone fragility across numerous diseases [11], including breast and prostate bone metastases [7, 12-14]). Yet, how tumor-derived factors reciprocally impair OCy mechanosensing under dynamic physical forces remains poorly understood. Data from our lab suggests that BCa cells shift the balance of downstream bone remodeling signals toward a pro-resorptive phenotype through their secreted signaling factors, even when subjected to anabolic fluid shear stresses [15, 16]. Furthermore, current standard-of-care treatments rely heavily on antiresorptives (e.g., bisphosphonates, RANKL inhibitors). While these target the remodeling secretome, they act strictly as palliatives and do not restore intrinsic OCy mechanosensitivity or restore lost bone. Therefore, there is a need to elucidate how tumor-specific factors disrupt Ocy mechanotransduction, both to understand why exercise and physical loading lose their protective efficacy and to uncover novel pathways for rescuing OCy function in metastatic bone disease To isolate tumor-induced disruptions to OCy mechanosensing, New Approach Methodologies (NAMs) utilize engineered 3D platforms that recapitulate key biochemical and physical aspects of the bone microenvironment. We utilize an engineered bone-mimetic scaffold combined with a multi-channel forced perfusion bioreactor. This polymeric and hydroxyapatite-coated scaffold recapitulates key biochemical features of the bone microenvironment, and supports osteogenic cell [16-19] and cancer [5, 20-22] cell function. It provides consistent pore size and distribution [23] and applies computationally modeled fluid shear stresses to standardize mechanical inputs [24-27]. Previously, using early-stage OCys (MLO-Y4s) in our platform, we showed that BCa-conditioned media impaired the canonical OCy mechanoresponse, preventing flow-induced downregulation of *rankl:opg* expression [16]. However, early-stage OCys lack key features of terminally differentiated cells, particularly sclerostin expression. Here, to overcome this limitation, we have extended our 3D platform to support mature OCys differentiated from a new cell line, Mesenchymal Progenitor Cells (MPC-2s), which form extensive dendritic networks, express sclerostin, and are sensitive to hormonal stimuli [28].

Although BCa cells are known to induce OCy senescence [12], loss of lacunar network integrity [14], and apoptosis [29], the impact of BCa molecular heterogeneity on OCy mechanosensing remains uncharacterized. Clinical BCa is categorized into four main clinical subtypes based on expression of hormone receptor (HR) and human epidermal growth factor receptor 2 (HER2): HR+/HER2−, HR+/HER2+, HR−/HER+, and triple negative (TN) [30]. These subtypes differ in patient prognosis, progression-free survival, and therapeutic response [31-33]. Most preclinical models rely on aggressive, reliably osteolytic TN breast cancer cell lines, despite the majority of metastatic patients presenting with ER+/HR+ disease. Furthermore, we have shown that fluid shear stresses within the bone microenvironment can dynamically alter the tumor secretome itself [34]. For example, fluid flow may paradoxically upregulate pro-resorptive and mechanosensitive factors in TN breast cancer cells, which may actively counteract the osteoprotective benefits of physical loading. Therefore, in this study, we investigated whether BCa-induced impairment of OCy mechanosensing is BCa subtype-dependent. Specifically, we evaluated mature OCy mechanoresponses under fluid shear stress when exposed to secretomes from ER+ versus TN cells. We also tested the downstream impacts of radiation, a frontline therapy for bone metastasis [35], on BCa-derived factors and their subsequent effects on the OCy mechanoresponse.

## MATERIAL & METHODS

### Breast Cancer Cell Culture

Two human breast cancer cell lines were used to generate conditioned media. Estrogen positive (ER+) breast cancer cells (ZR-75-1, ATCC) were cultured in standard culture conditions (37°C, 5% CO_2_) in complete media consisting of RPMI-1640 (ATCC Ref: 30-2001) supplemented with 10% non-heat-inactived Fetal Bovine Serum (FBS) (Avantor: 89510-186), and 1% Penicillin-Streptomycin (P-S, Sigma Aldrich: P4333). Triple negative (TN) breast cancer cells (MDA-MB231, ATCC) were cultured in standard culture conditions in complete media consisting of Dulbecco’s Modified Eagle Medium (DMEM, Sigma Aldrich: M0644) supplemented with 10% FBS (R&D Systems: S11550H) and 1% P-S. Media was replenished 2-3 times per week.

### Breast Cancer Cell Irradiation and Generation of Conditioned Media (CM)

When ZR-75-1 and MDA-MB-231 cultures reached 70% confluency, complete media was replaced with low serum media. Cells were immediately irradiated inside a Faxitron Cabinet X-ray System Model RX-650 (Lincolnshire, IL) at 115 kv and 5.33 rad/s to deliver doses of either 0 Gy (control) or 2 Gy. Following irradiation, cells were then cultured for an additional 24 hours, their conditioned media (CM) collected and centrifuged at 1000 RPM for 10 minutes. The resulting supernatant was stored at -80°C in aliquots to avoid multiple freeze/thaw cycles.

### Mesenchymal Progenitor Cell Culture

Mesenchymal Progenitor Cells (‘MPCs’) were expanded up to passage 17 in an undifferentiated state at 33°C and 5% CO_2_ in Iscove’s Modified Dulbecco’s Medium (IMDM) supplemented with L-glutamine (Gibco Ref: 12440053), 10% FBS and 1% P-S. Following expansion, MPCs were seeded into scaffolds (see §*Bone-mimicking Scaffold and MPC Seeding*), and cultured osteogenic induction media [Minimum Essential Medium α (MEM-α) with L-glutamine (Millipore Ref: M4526) supplemented with 10% FBS, 1% P-S, ascorbic acid (50 μg/ml) and β-glycerophosphate (10 mM) in standard culture conditions. Osteogenic media was replaced every 2-3 days.

### Bone-mimicking Scaffold and MPC Seeding

Bone-mimetic scaffolds were fabricated with poly(lactide-co-glycolide) (PLGA) and hydroxyapatite (HA) particles using a gas foaming-particulate leaching method. Briefly, 8 mg of HA particles (Sigma-Aldrich, 677418), 8 mg of poly(lactide-coglycolide) microspheres, and 152 mg of NaCl crystals (sieved to a range of 250–400 μm in diameter), were pressure molded (Carver Press) into disks (8mm diameter, ∼2mm height) [19-21]. Disks were then subjected to a gas-foaming/particulate leaching technique to create a mineralized surface. Prior to cell-seeding, scaffolds were disinfected in 70% ethanol for 30 min, washed 4× with sterile PBS, and then seeded dropwise with 500,000 MPCs per scaffold. Seeded scaffolds were maintained on an orbital shaker throughout all experiments. Following a 48 hr attachment period, osteogenic media was introduced and maintained for either 2 or 3 weeks to induce differentiation.

### Forced Perfusion Bioreactor

For perfusion optimization experiments, following 2 or 3 weeks of differentiation (2 Wks, 3 Wks), dynamice forced perfusion was applied to MPC-laden scaffolds for 3 consecutive days via our custom multi-channel bioreactor. An inlet volumetric flow rate of 5 mL/min, corresponding to ∼60 mPa median interstitial wall shear stresses [27], was delivered at 1 Hz for either 1 or 2 hrs/day. Static groups served as controls. Following the final perfusion bout, scaffolds were immediately processed for downstream analyses. For the tumor conditioned media experiments, following 2 weeks of differentiation, MPC-laden scaffolds were incubated for 24 hrs with breast cancer cell CM diluted 1:1 in osteogenic media. Then, scaffolds were subjected to fluid flow (5 mL/min at 1Hz) for 2 hrs per day for 3 consecutive days. Constructs were processed for analysis immediately following the final loading bout.

### Parathyroid Hormone Stimulation

Following 2 weeks of differentiation, MPC-seeded scaffolds received 0 or 50 nM of human parathyroid hormone (PTH 1-34, Sigma-Aldrich P3796) in osteogenic media. After 48 hrs, scaffolds were processed for downstream analysis.

### Histology for MPC Morphology

Scaffolds were fixed with 4% paraformaldehyde for 30 min, and stored in PBS prior to whole-mount histochemical staining. Cells within scaffolds were permeabilized with 0.05% (v/v) Triton-X and incubated with DAPI (1:1000, Sigma-Aldrich, D9542) and phalloidin (1:1000, Thermo Fisher Scientific A12379 or Cayman Chemical Company 20549) for 15 minutes at room temperature. Fluorescent images were acquired using a confocal microscope (Nikon A1R) and visualized (Fiji ImageJ) for evaluation of osteocyte morphology.

### MPC Gene Expression

Total RNA was extracted using TRIzol reagent under RNase-free conditions and reverse-transcribed into cDNA (Qiagen, 205311). Real-time quantitative PCR (qPCR) was performed using a Rotor-Gene Q system (Qiagen, 9001862) with QuantiNova SYBR Green PCR master mix (Qiagen: 208252) and custom-designed primers (Suppl. Table S1). Gene expression levels for *Sost* (sclerostin), *Tnfsf11* (RANKL), and *Tnfrsf11b* (OPG) were determined using the efficiency-corrected 2^-ΔΔCT^) method. Target gene expression was normalized to *Hprt* (hypoxanthine phosphoribosyltransferase),which we determined to be invariant across all treatment conditons.

### Antibody Array with Breast Cancer CM

Protein expression profiles in breast cancer cell CM were characterized using a membrane antibody array targeting 84 human oncology-related proteins (Proteome Profiler Human XL Oncology Array, R&D Systems: ARY026). Aliquoted CM supernatants were diluted and incubated with the array membranes overnight according to the manufacturer’s instructions. Following wash steps to remove unbound proteins, membranes were incubated with biotinylated detection antibodies and streptavidin-HRP. Chemiluminescent signals were captured using an Azure Imaging System (Azure Biosystems) with a fixed 10 min exposure time. Densitometric analysis was conducted in MATLAB by measuring the integrated pixel density of each capture spot, followed by background subtraction and normalization to internal reference spots.

### Statistical Analysis

All experiments were performed with 3–5 independent biological replicates, each containing 2–4 technical replicates per outcome measure. Data points represent the mean value of technical replicates for individual biological replicates. For perfusion experiments, loaded groups within each timepoint (2 Wks vs. 3 Wks) or loading duration (1 h vs. 2 h) were normalized to their respective static controls and evaluated using Student’s t-test. Differences between PTH treatments (0 vs. 50 nM) were evaluated via Student’s t-test. Main effects and interactions of BCa cell subtype (ER+ vs. TN) and irradiation status (0 vs. 2 Gy) were evaluated using a two-way full-factorial ANOVA followed by Bonferroni post-hoc tests for multiple comparisons. Outliers were detected and excluded using the Interquartile Range (IQR) / Quantile Range method. All data are represented as the mean ± SEM across all biological replicates. Statistical significance was set at α *<* 0.05.

## RESULTS

### 1. MPCs cells in 3D form interconnecting dendrites following 2 weeks of osteogenic differentiation

To establish a physiologically relevant 3D model of mature OCy, we verified the capacity of MPCs to differentiate and form interconnected dendritic networks within mineralized scaffolds (Fig1). Following two weeks of osteogenic induction, whole-mount confocal microscopy revealed robust cellular attachment and the development of extensive dendritic networks across the scaffold surface (Fig1B). Differentiated MPCs adopted a characteristic osteocytic morphology, exhibiting numerous elongated dendritic processes that formed functional intercellular connections with neighboring cells (Fig1C). Together, these findings demonstrate that our 3D engineered scaffold system successfully supports MPC differentiation into mature, dendrite-bearing OCys capable of forming an interconnected network.

**Figure 1:**
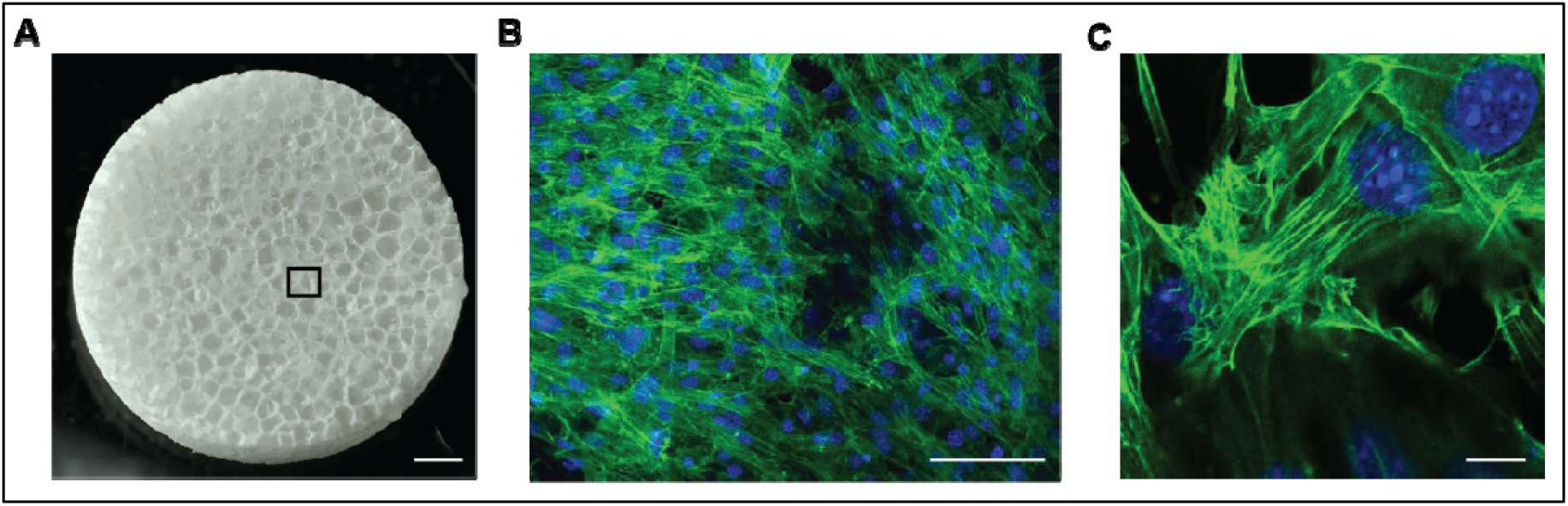
Osteogenic MPC cells in 3D form interconnecting dendrites by 2 weeks of osteogenic differentiation. Following 2 weeks of osteogenic differentiation within bone-mimetic scaffolds, dendrite formation and morphology of MPCs were assessed via F-actin (green, phalloidin) and nuclear (blue, dapi) staining. **(A)** Porous, hydroxyapatite coated bone mimetic scaffold. Scale bar = 1 mm. **(B)** Osteogenic MPCs formed a network of interconnected dendrites. Representative confocal image of osteogenic MPCs on the surface of scaffolds via whole mount histology. Scale bar = 100 µm. **(C)** High-resolution confocal imaging shows multiple dendritic connection between neighboring osteogenic MPCs. Scale bar = 10 µm.

### 2. Osteogenic MPC cells in 3D exhibit the appropriate mechanoresponse to fluid shear and hormone stimulation

To optimize the differentiation duration and canonical mechanoresponse to fluid shear stress (FSS) protocol, MPC-laded scaffolds were evaluated across different time points and daily perfusion durations. Following either 2 or 3 weeks of osteogenic differentiation, a significant, flow-induced downregulation of *Sost* expression was observed after 2 weeks of differentiation, whereas scaffolds differentiated for 3 weeks showed no significant Sost response to flow (Fig2A). Although expression of RANKL:OPG (*Tnfsf11:Tnfrsf11b*) decreased after flow at both time points (Fig2B), this shift was primarily driven by reductions in OPG (*Tnfrsf11b*) at 2 weeks (Fig2D) rather than consistent changes in RANKL (*Tnfsf11*) (Fig2C). Based on the significant *Sost* suppression observed at 2 weeks, this differentiation duration was selected to evaluate the impact of daily flow duration (1 h vs. 2 hr/day for 3 days) (Fig3). Both 1-hour and 2-hour daily FSS treatments successfully suppressed *Sost* and reduced RANKL:OPG (*Tnfsf11:Tnfrsf11b*) compared to static controls (Figs 3A, 3B).

**Figure 2:**
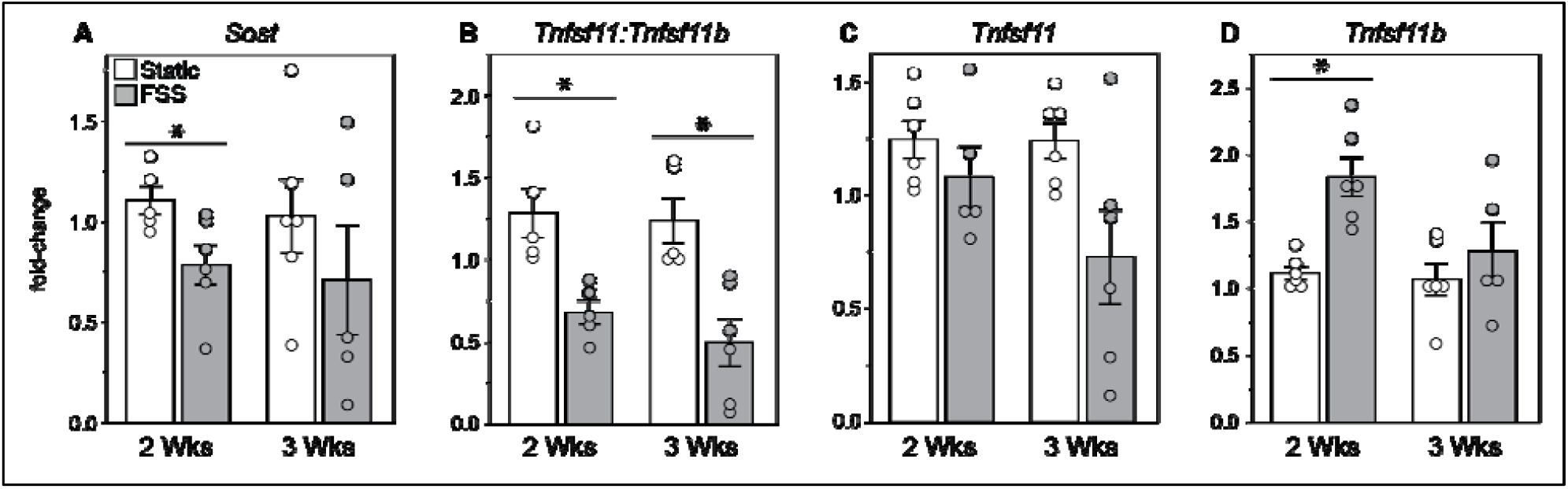
Osteogenic MPCs in 3D exhibit a stronger mechanoresponse to fluid shear after 2 weeks of differentiation. Following either 2 (2 Wks) or 3 (3 Wks) weeks of osteogenic differentiation within bone mimetic scaffolds, MPCs were subjected to 1 hr per day of unsteady (1 Hz) forced fluid shear stress (FSS) for 3 days. **(A)** Genetic expression of *Sost* decreased after 2 Wks, but not 3 Wks. **(B)** *Tnfsf11:Tnfsf11b* (RANKL:OPG) decreased following 3 days of FSS at both differentiation lengths. **(C)** *Tnfsf11* (RANKL) displayed an inconsistent response to FSS. **(D)** Shear-induced reductions in *Tnfsf11b* (OPG) therefore contributed the most to overall reductions in *Tnfsf11:Tnfsf11b* (RANKL:OPG) after 2 Wks of differentiation. Error bars = SEM. *p<0.05, t-test Static vs. FSS.

**Figure 3:**
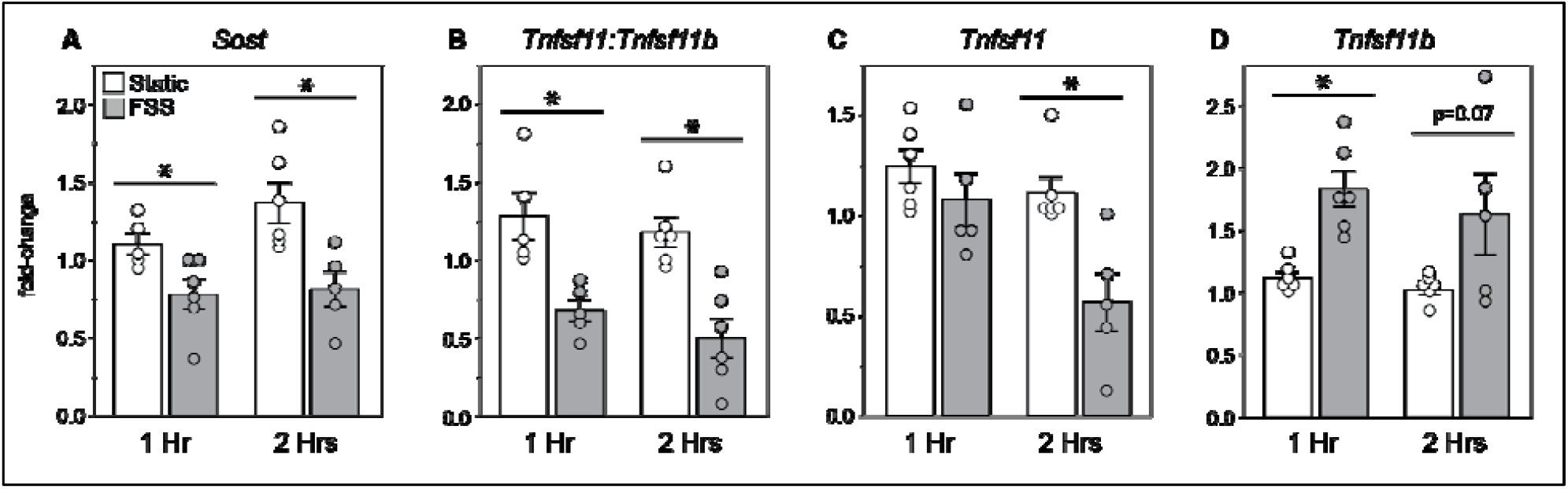
Osteogenic MPCs in 3D exhibit a more consistent mechanoresponse to fluid shear stress after 2 weeks of differentiation. Following 2 weeks of osteogenic differentiation within bone-mimetic scaffolds, MPCs were subjected to either 1 (1 Hr) or 2 (2 Hr) hrs per day of unsteady (1 Hz) forced fluid shear stress (FSS) for 3 days. Expression of both **(A)** *Sost* and **(B)** *Tnfsf11:Tnfsf11b* (RANKL:OPG) decreased following 1 Hr and 2 Hrs of FSS. **(C)** However, *Tnfsf11* (RANKL) significantly decreased only after 2 Hrs of FSS, while **(D)** *Tnfsf11b* (OPG) significantly increased only after 1 Hr of FSS with a trend after 2 Hrs. Note that the 1 Hr data is also displayed in Fig2. Error bars = SEM. *p<0.05, t-test Static vs. FSS.

Notably, increasing daily FSS duration to 2 hr/day elicited a more consistent anti-catabolic response, driving a significant reduction in RANKL (*Tnfsf11*) expression (Fig3C), while 1 hour of FSS primarily elevated OPG (*Tnfrsf11b*) expression (Fig3D). Together, these parameter optimization experiments establish that 2 weeks of osteogenic differentiation paired with 2 hr/day of dynamic perfusion yields the most consistent canonical OCy mechanoresponse in our 3D platform.

To confirm that 3D mature MPC-derived OCy maintain physiological endocrine sensitivity in addition to mechanosensitivity, 2-week differentiated scaffolds were treated with exogenous parathyroid hormone (50 nM PTH 1-34 for 48 hr) (Fig4). PTH treatment significantly upregulated RANKL (*Tnfsf11*) gene expression (Fig4C) and showed a trend toward increased OPG (*Tnfrsf11b*) expression (Fig4D), resulting in a trend toward an elevated RANKL:OPG (*Tnfsf11:Tnfrsf11b*) (Fig4B), characteristic of PTH-mediated bone remodeling signaling. In contrast, *Sost* expression remained unchanged following 48 hr of continuous PTH exposure (Fig4A). These findings demonstrate that 3D MPC-derived OCy retain intact, classical receptor-mediated responses to systemic hormonal cues.

**Figure 4:**
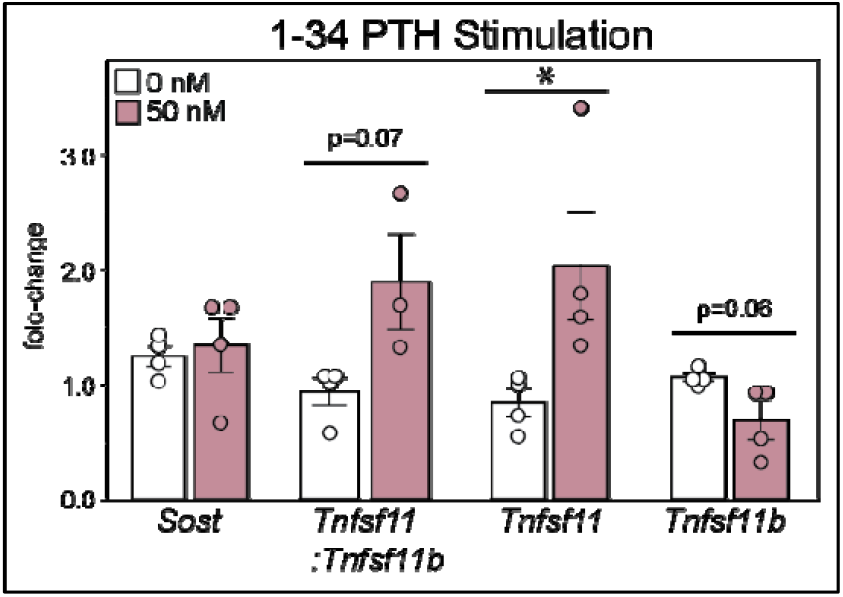
Osteogenic MPCs in 3D increase gene expression of RANKL following parathyroid hormone (PTH) treatment. Following 2 weeks of osteogenic differentiation within bone-mimetic scaffolds, MPCs were stimulated with 50 nM 1-34 PTH for 48 hrs. There was a trend for increased expression of *Tnfsf11:Tnfsf11b* (RANKL:OPG) due to significant increases in *Tnfsf11* (RANKL) and a trend for increased *Tnfsf11b* (OPG). No significant changes occurred in *Sost* expression. Error bars = SEM. *p<0.05, t-test 0 vs. 50 nM.

### 4. Breast cancer subtype and irradiation status differentially disrupt osteogenic MPCs’ mechanoresponsiveness

To determine how BCa molecular subtype and radiation therapy alter tumor-induced OCy dysfunction, 3D mature MPC-derived OCys were exposed to conditioned media (CM) from estrogen receptor–positive (ER+; ZR-75-1) or triple-negative (TN; MDA-MB-231) BCa cells generated 24 hr post-irradiation (0 vs. 2 Gy), and subsequently subjected to fluid shear stress (FSS; 2 h/day for 3 days) (Fig5). The flow-induced suppression of *Sost* was differentially altered by BCa subtype and radiation status. Osteogenic MPCs cultured with non-irradiated (0 Gy) ER+ CM retained their canonical mechanoresponse, showing significant *Sost* downregulation following dynamic perfusion. However, this flow-induced reduction in *Sost* was completely abolished when MPCs were exposed to CM from irradiated (2 Gy) ER+ cells (Fig5A). In contrast, exposure to TN CM completely blunted flow-induced *Sost* suppression regardless of radiation status, with MPCs showing no significant change in *Sost* expression following flow in either the 0 Gy or 2 Gy TNBC CM groups (Fig5A).

**Figure 5:**
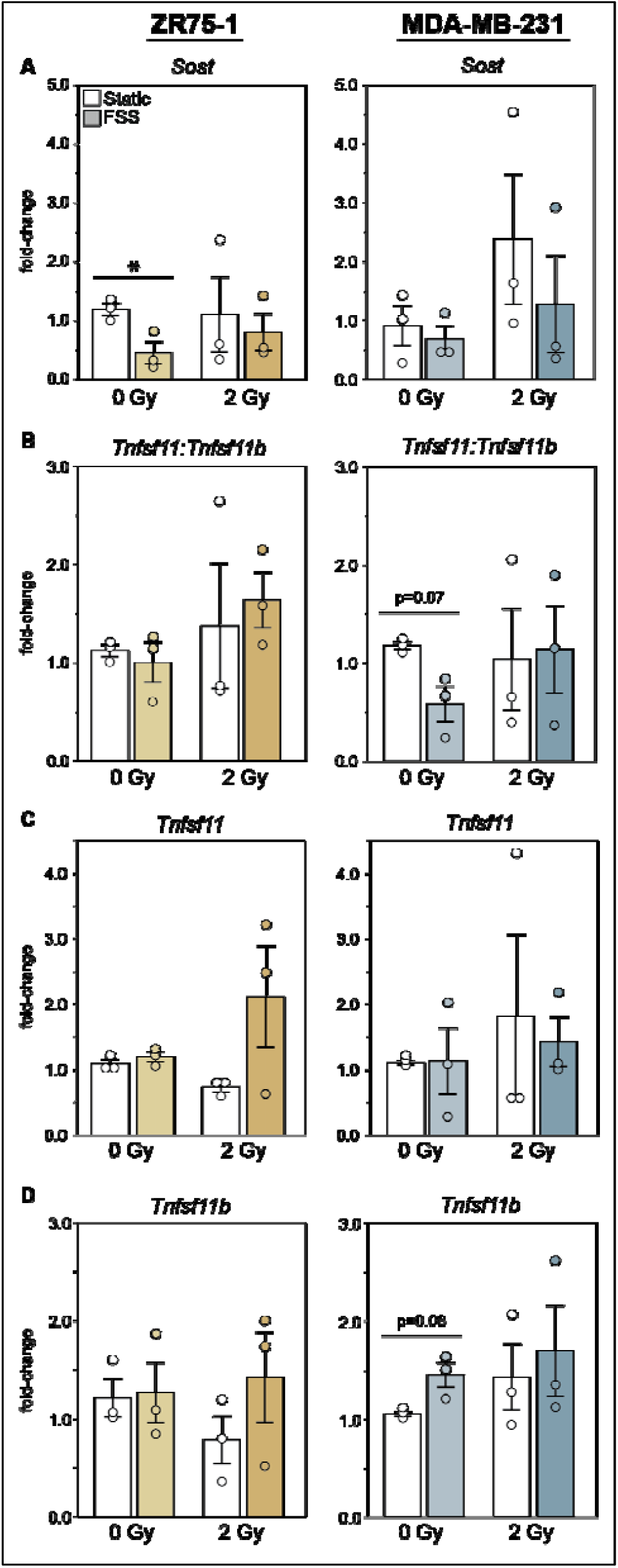
The breast cancer cell secretome impacts the mechanoresponse of osteogenic MPCs to fluid shear in a subtype-dependent manner. Following 2 weeks of osteogenic differentiation within bone-mimetic scaffolds, MPCs were cultured with conditioned media (CM) from BCa cells for 24 hours. CM was collected from Estrogen-Receptor+ (ER+, ZR75-1) or Triple Negative (TN, MDA-MB-2331) BCa cells 24 hrs following treatment with 0 or 2 Gy irradiation. After, MPCs were subjected to 2 hrs per day of unsteady (1 Hz) forced fluid shear stress (FSS) for 3 days. **(A)** Osteogenic MPCs retained their ability to reduce expression of *Sost* following FSS despite the presence of CM from ER+ cells. However, this response to FSS disappeared after ER+ cells received 2 Gy. MPCs did not reduce *Sost* expression when cultured with CM from TN cells irrespective of irradiation status. **(B)** *Tnfsf11:Tnfsf11b* (RANKL:OPG) expression in MPCs cultured with ER+ CM did not decrease following to FSS regardless of irradiation status. MPCs with TN CM exhibited a trend for decreased *Tnfsf11:Tnfsf11b* (RANKL:OPG) only with culture from 0 Gy cells. **(C)** *Tnfsf11* (RANKL) did not decrease following FSS in any CM condition. **(D)** A trend for increased *Tnfsf11b* (OPG) occurred only in MPCs treated with 0 Gy TNBC CM, driving the observed changes in *Tnfsf11:Tnfsf11b* (RANKL:OPG). Error bars = SEM. *p<0.05, 2-Way Full-factorial ANOVA [Factors: Loading status (Static vs. FSS), Radiation dose (0 vs. 2 Gy)].

Evaluation of expression of remodeling signals revealed that osteogenic MPCs exposed to ER+ CM failed to decrease RANKL:OPG (*Tnfsf11:Tnfrsf11b*) following FSS, irrespective of radiation dose (Fig5B). Conversely, MPCs cultured with 0 Gy TN CM displayed a trend toward a reduced RANKL:OPG post-flow (Fig5B), which was primarily driven by a trend toward increased OPG (*Tnfrsf11b*) expression (Fig5D) rather than changes in RANKL (*Tnfsf11*) (Fig5C). This flow-induced OPG shift was absent in MPCs treated with 2 Gy TN CM (Fig5D). Together, these results demonstrate that OCy mechanosensitivity is differentially compromised by BCa molecular subtype, and that low-dose radiation may further exacerbates secretome-mediated mechanosensory impairment in ER+ disease.

### 5. Profiling of the breast cancer cell secretome displayed subtype-dominant and radiation-sensitive alterations

To identify the tumor-derived soluble factors mediating these subtype- and radiation-dependent impacts on osteocytes, we profiled 84 oncology-related proteins in ER+ and TN CM 24 hr post-irradiation using membrane antibody arrays. Comparative baseline analysis revealed that molecular subtype was the primary driver of tumor secretome composition (Fig6). Down-selection for the top 15 most abundant proteins consistently detected across biological replicates (n=3) demonstrated distinct proteomic profiles between subtypes. While both lines secreted Cathepsin D and ENPP-2/Autotaxin at high levels, overall abundance was substantially higher in ER+ CM. Furthermore, the top secreted proteins in ER+ CM were largely stable following 2 Gy irradiation, with ErbB2 and ErbB3/HER3 consistently ranking among the highest expressed factors across 5 of 6 replicates. In contrast, the high-abundance profile of TN CM exhibited greater sensitivity to radiation, with only Axl, Dkk-1, and Amphiregulin consistently remaining among the most abundant proteins regardless of irradiation status.

**Figure 6:**
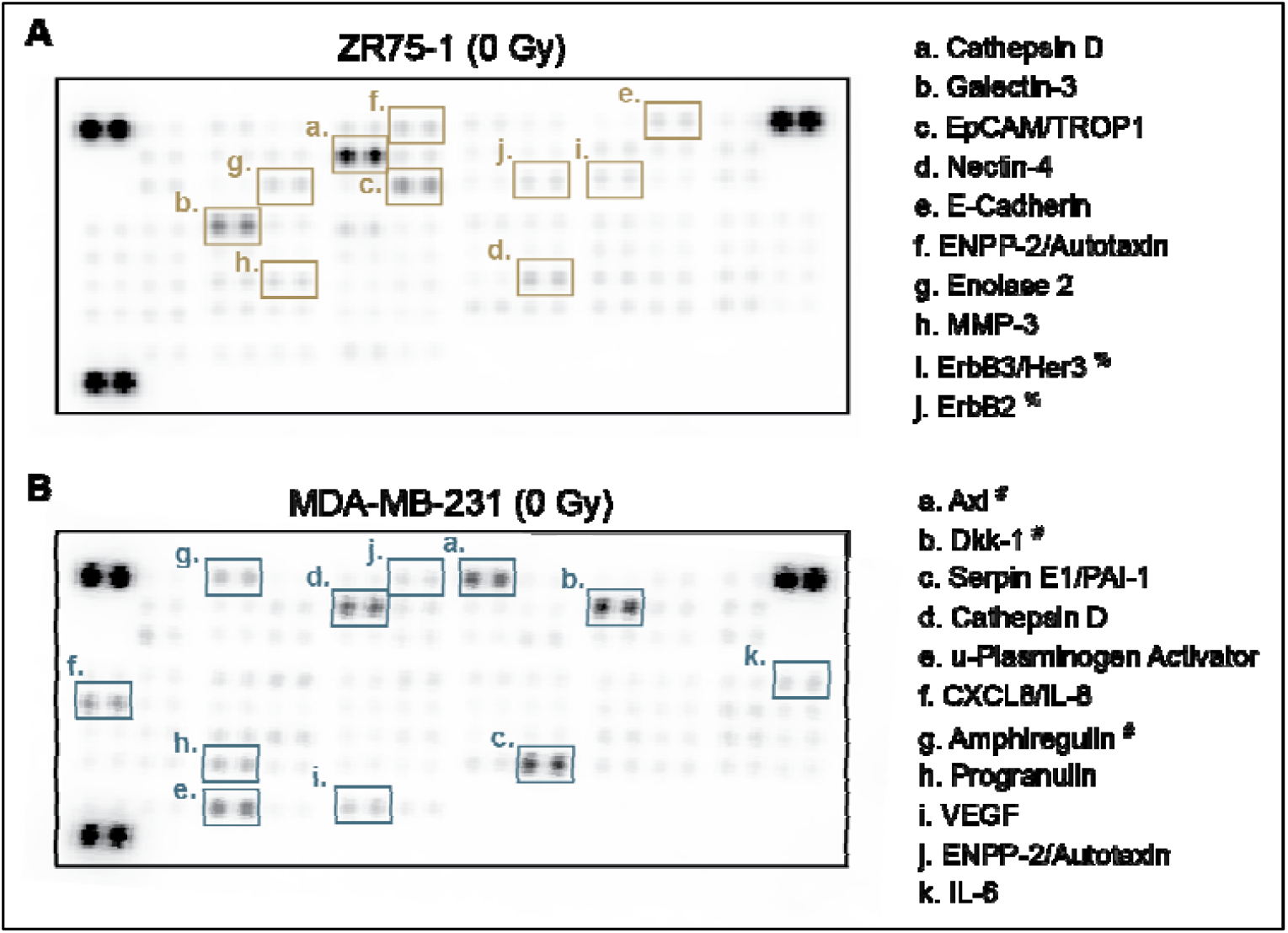
Breast cancer cell secretome substantially differed between subtypes. Conditioned media Estrogen-Receptor+ (ER+, ZR75-1) or Triple Negative (TN, MDA-MB-2331) BCa cells was collected 24 hours after irradiation. Representative antibody arrays from **(A)** ER+ (0 Gy) and **(B)** TN BCa cells (0 Gy). %Proteins that did not consistently remain at high abundance following 2 Gy radiation. #Proteins that remained consistently high in abundance, independent of radiation.

To evaluate specific radiation-induced secretome shifts, proteins were filtered for consistent directionality (increased or decreased post-radiation) and low variance (coefficient of variation ≤30%) across biological replicates. Radiation-induced changes were generally modest and displayed higher inter-replicate variability compared to baseline subtype differences. Nevertheless, directional trends emerged between subtypes. Low-dose radiation (2 Gy) tended to induce an overall upregulation of protein secretion in ER+ cells, whereas it primarily downregulated protein secretion in TN cells (Fig7, Table 1). These subtype-specific secretome profiles highlight key candidate mediators, such as Dkk-1, Cathepsin D, and ErbB family receptors, that may drive the differential loss of OCy mechanosensitivity.

**Figure 7:**
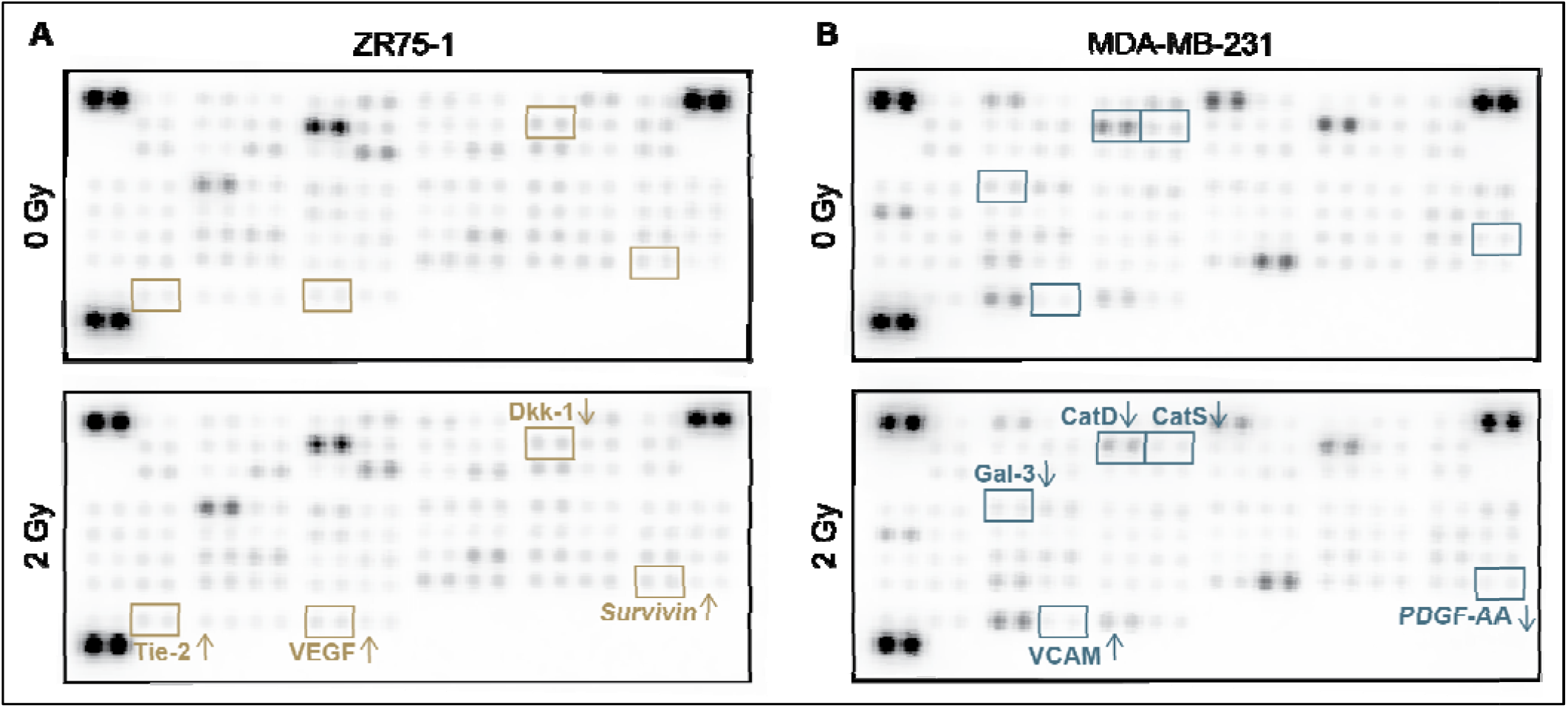
The breast cancer cell secretome was mostly insensitive to radiation. Representative antibody arrays from **(A)** ER+ breast cancer cells and **(B)** TNBC breast cancer cells reveal modest changes due to radiation with high variability. Interestingly, among proteins down-selected (details?), radiation tended to cause increases in protein secretion from ZR75-1 (ER+) cells and decreases in proteins from MDA-MD-231 (TNBCA) cells.

**Table 1:**
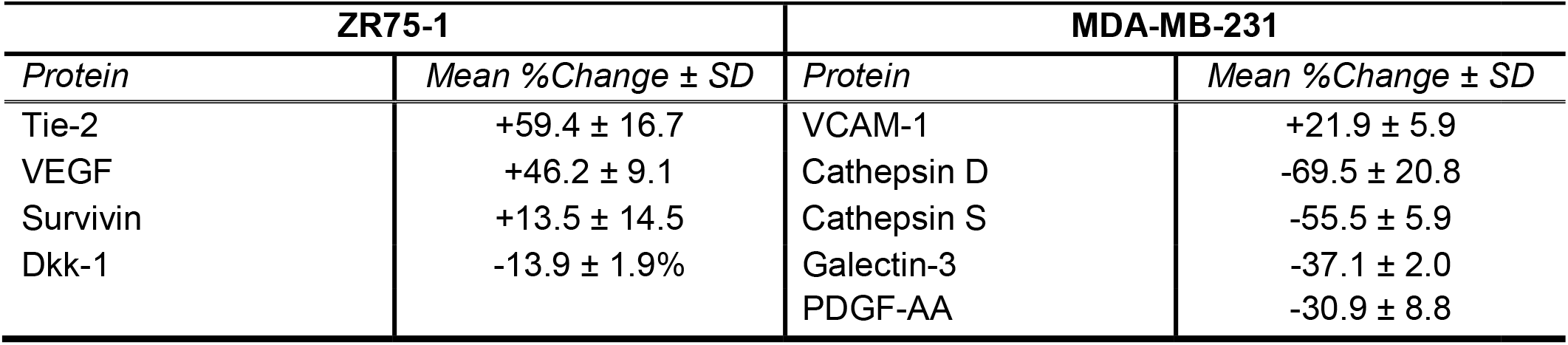
Notable Changes in Proteins Within the Breast Cancer Cell Secretome Due to 2 Gy Radiation.

## DISCUSSION

In this study, we established a physiologically relevant 3D bone-mimetic platform to interrogate how breast cancer (BCa) molecular heterogeneity and frontline radiation therapy modulate tumor-induced osteocyte (OCy) mechanosensory dysfunction. By extending our forced perfusion model to support mature, dendrite-bearing MPC-2 osteocytes capable of expressing sclerostin and maintaining canonical mechanoresponsiveness, we isolated the contribution of the tumor secretome to impaired skeletal mechanotransduction. Our primary finding is that BCa-induced disruption of OCy mechanosensing is subtype-dependent. While non-irradiated ER+ (ZR-75-1) secretomes preserved flow-induced *Sost* suppression, triple-negative (TN; MDA-MB-231) secretomes abolished canonical OCy mechanoresponsiveness regardless of loading status. Also, low-dose (2 Gy) radiation eliminated the protective mechanoresponse in ER+ cultures, demonstrating that therapeutic interventions can dynamically alter tumor secretomes to exacerbate OCy dysfunction. Proteomic profiling confirmed that molecular subtype is the dominant driver of the BCa secretome, highlighting key candidate mediators (e.g, Cathepsin D) that may act as paracrine drivers of tumor-induced bone fragility.

MPCs in our 3D scaffold reached maturity producing dendrites and sclerostin. Currently, only a few cell lines exist that reliably result in mature, sclerostin-expressing OCys, such as the IDG-SW3 [36] and OCY454 [37] cell lines. While these cells have been used to create 3D models (e.g., [38-40]), only a few incorporated dynamic loading. Rindt et. al. developed a 3D system in which IDG-SW3 osteocytes cultured in mineralized collagen scaffolds reduced sclerostin and RANKL:OPG in response to applied perfusion and compression, similar to our results [41]. Likewise, IDG-SW3 osteocytes cultured in a combined cartilage-bone bilayer hydrogel also reduced sclerostin and RANKL:OPG following compression [42]. Both studies reported other aspects of the mature OCy mechanoresponse, such as upregulation of PGE2. Overall, these types of 3D systems are necessary to study OCY mechanobiology in a more physiologically relevant microenvironment.

A central advance of our study is demonstrating that tumor-induced OCy mechanosensory impairment is dependent upon BCa molecular subtype. To date, the majority of preclinical bone oncology models rely on aggressive, osteolytic TNBC lines (e.g., MDA-MB-231). But, over 70% of clinical BCa bone metastases originate from ER+/HR+ primary tumors [43]. Our findings highlight a physiological contrast between these subtypes: while ER+ (ZR-75-1) secretomes permitted baseline flow-induced *Sost* suppression, TN secretomes abolished canonical OCy mechanoresponsiveness. This differential sensitivity mirrors clinical observations where HR+ bone metastatic patients often maintain better structural bone health and respond more favorably to physical activity compared to those with TN [44]. While our work established this benchmark by comparing the most clinically prevalent subtype (ER+) against the most widely studied model (TNBC), future work should incorporate the full clinical spectrum of BCa subtypes, including HER2-overexpressing and Luminal B lines, to map how diverse oncogenic drivers uniquely dysregulate OCy mechanosensing.

Our proteomic array profiling provided key mechanistic insights into the paracrine factors driving these subtype-specific responses. Dkk-1 was consistently, and selectively, secreted by TN BCa cells, in line with other reports [45]. Dkk-1 is a potent inhibitor of canonical Wnt/β-catenin signaling, and may compensate for reduced sclerostin [46]. In addition to Dkk-1, TN secretomes contained elevated levels of pro-inflammatory cytokines such as IL-6 and IL-8. Chronic exposure to inflammatory cytokines disrupts focal adhesion maturation and alters cytoskeletal stiffness in osteocytes, which may desensitize the mechanotransduction machinery required for fluid flow sensing. Conversely, ER+ secretomes contained distinct factors such as Cathepsin D, ENPP-2/Autotaxin, and ErbB receptors. To fully resolve these complex paracrine networks, future studies should deploy unbiased liquid chromatography–tandem mass spectrometry (LC-MS/MS) discovery proteomics to capture the complete secretome profile under dynamic mechanical loading.

Additionally, future work should determine whether any of identified candidates, such as Dkk-1, Autotaxin, or radiation-induced inflammatory factors, to drive OCy mechanosensory failure.

Our data suggests that radiation therapy, a standard palliative treatment for bone metastases, alters the BCa secretome with implications for bone tissue mechanosensitivity. Low-dose radiation (2 Gy) abolished the flow-induced *Sost* suppression previously preserved in ER+ co-cultures, consistent with reported negative effects of radiation on bone [47]. In contrast, high-magnitude, low-frequency vibration, applied after 8 Gy of radiation, recovered aspects of OCy function [48], or mitigated tumor-induced bone loss [9]. Determining the combined effects of loading and radiation, and their sequence [34], is important for successful incorporation of physical therapy or clinical vibration of metastasis patients.

## ACKNOWLEDGEMENTS

The authors acknowledge: Dr. Tin Tin Su for allowing them to use her irradiator in this study; Dr. Sarah Calve and Terren Geohring for technical support on scaffold histology; and Dr. Jian Tay for technical support on antibody array image analysis.

## FUNDING

The authors gratefully acknowledge funding from the following sources: NSF BMMB-2047187

T32AR080630

Anschutz-Boulder Nexus Award #9866

## AUTHOR CONTRIBUTIONS

Conceptualization: WRT, MEL

Data Curation: ASA, CB, EG, ENB, AMB, TKR

Analysis: ASA, CB, ENB, TKR, MEL

Funding Acquisition: MEL

Methodology: ASA, CB, EG, WRT, MEL

Writing – original draft: ASA, CB, MEL

Writing – review & editing: All authors

## REFERENCES

1. Robling, A.G. and L.F. Bonewald, The Osteocyte: New Insights. Annu Rev Physiol, 2020. 82: p. 485–506.

2. Schaffler, M.B., et al., Osteocytes: master orchestrators of bone. Calcif Tissue Int, 2014. 94(1): p. 5–24.

3. Fan, Y., et al., Skeletal loading regulates breast cancer-associated osteolysis in a loading intensity-dependent fashion. Bone Res, 2020. 8: p. 9.

4. Huang, M., et al., Mechanical loading attenuates breast cancer-associated bone metastasis in obese mice by regulating the bone marrow microenvironment. J Cell Physiol, 2021.

5. Lynch, M.E., et al., In vivo tibial compression decreases osteolysis and tumor formation in a human metastatic breast cancer model. J Bone Miner Res, 2013. 28(11): p. 2357–67.

6. Rummler, M., et al., Mechanical loading prevents bone destruction and exerts anti-tumor effects in the MO PC315.BM. Luc model of myeloma bone disease. Acta Biomater, 2021. 119: p. 247–258.

7. Wang, S., et al., Moderate tibial loading and treadmill running, but not overloading, protect adult murine bone from destruction by metastasized breast cancer. Bone, 2021: p. 116100.

8. Ziouti, F., et al., Prevention of Bone Destruction by Mechanical Loading Is Not Enhanced by the Bruton’s Tyrosine Kinase Inhibitor CC-292 in Myeloma Bone Disease. Int J Mol Sci, 2021. 22(8).

9. Pagnotti, G.M., et al., Low intensity vibration mitigates tumor progression and protects bone quantity and quality in a murine model of myeloma. Bone, 2016. 90: p. 69–79.

10. Wang, S., et al., Moderate tibial loading and treadmill running, but not overloading, protect adult murine bone from destruction by metastasized breast cancer. Bone, 2021. 153: p. 116100.

11. Kitase, Y. and M. Prideaux, Regulation of the Osteocyte Secretome with Aging and Disease. Calcif Tissue Int, 2023. 113(1): p. 48–67.

12. Kaur, J., et al., Single-Cell Transcriptomic Analysis Identifies Senescent Osteocytes That Trigger Bone Destruction in Breast Cancer Metastasis. Cancer Res, 2024. 84(23): p. 3936–3952.

13. Liu, S., et al., Overexpression of Lrp5 enhanced the anti-breast cancer effects of osteocytes in bone. Bone Res, 2021. 9(1): p. 32.

14. Hemmatian, H., et al., Reorganization of the osteocyte lacuno-canalicular network characteristics in tumor sites of an immunocompetent murine model of osteotropic cancers. Bone, 2021. 152: p. 116074.

15. Wang, W., et al., Mechanically-Loaded Breast Cancer Cells Modify Osteocyte Mechanosensitivity by Secreting Factors That Increase Osteocyte Dendrite Formation and Downstream Resorption. Front Endocrinol (Lausanne), 2018. 9: p. 352.

16. Sarazin, B.A., et al., Bone-homing metastatic breast cancer cells impair osteocytes’ mechanoresponse in a 3D loading model. Heliyon, 2023. 9(10): p. e20248.

17. Davis, H.E., et al., Osteogenic response to BMP-2 of hMSCs grown on apatite-coated scaffolds. Biotechnol Bioeng, 2011. 108(11): p. 2727–35.

18. Davis, H.E., et al., Biomimetic scaffolds fabricated from apatite-coated polymer microspheres. J Biomed Mater Res A, 2009. 90(4): p. 1021–31.

19. Lynch, M.E., et al., Three-Dimensional Mechanical Loading Modulates the Osteogenic Response of Mesenchymal Stem Cells to Tumor-Derived Soluble Signals. Tissue Eng Part A, 2016. 22(15-16): p. 1006–15.

20. Pathi, S.P., et al., A novel 3-D mineralized tumor model to study breast cancer bone metastasis. PLoS One, 2010. 5(1): p. e8849.

21. Pathi, S.P., et al., Hydroxyapatite nanoparticle-containing scaffolds for the study of breast cancer bone metastasis. Biomaterials, 2011. 32(22): p. 5112–22.

22. Fischbach, C., et al., Engineering tumors with 3D scaffolds. Nat Methods, 2007. 4(10): p. 855–60.

23. Harris, L.D., B.S. Kim, and D.J. Mooney, Open pore biodegradable matrices formed with gas foaming. J Biomed Mater Res, 1998. 42(3): p. 396–402.

24. Liu, B., et al., Perfusion applied to a 3D model of bone metastasis results in uniformly dispersed mechanical stimuli. Biotechnol Bioeng, 2018. 115(4): p. 1076–1085.

25. Liu, B., et al., Multiphysics simulation of a compression-perfusion combined bioreactor to predict the mechanical microenvironment during bone metastatic breast cancer loading experiments. Biotechnol Bioeng, 2021. 118(5): p. 1779–1792.

26. Han, S., et al., Flow inside a bone scaffold: Visualization using 3D phase contrast MRI and comparison with numerical simulations. J Biomech, 2021. 126: p. 110625.

27. Venkatesh, S., et al., High-fidelity computational fluid dynamics modeling to simulate perfusion through a bone-mimicking scaffold. Comput Biol Med, 2025. 186: p. 109637.

28. Prideaux, M., et al., Generation of two multipotent mesenchymal progenitor cell lines capable of osteogenic, mature osteocyte, adipogenic, and chondrogenic differentiation. Sci Rep, 2021. 11(1): p. 22593.

29. Wasi, M., et al., Different effects of moderate tibial loading and Yoda1 on breast cancer-induced osteolysis in aged mice. Bone, 2025. 197: p. 117517.

30. Perou, C.M., et al., Molecular portraits of human breast tumours. Nature, 2000. 406(6797): p. 747–52.

31. Rouzier, R., et al., Breast cancer molecular subtypes respond differently to preoperative chemotherapy. Clin Cancer Res, 2005. 11(16): p. 5678–85.

32. Sorlie, T., et al., Gene expression patterns of breast carcinomas distinguish tumor subclasses with clinical implications. Proc Natl Acad Sci U S A, 2001. 98(19): p. 10869–74.

33. Sorlie, T., et al., Repeated observation of breast tumor subtypes in independent gene expression data sets. Proc Natl Acad Sci U S A, 2003. 100(14): p. 8418–23.

34. Venkatesh, S., et al., Radiation Modulates the Mechanoresponse of Bone-Homing Triple-Negative Breast Cancer Cells. Tissue Eng Part A, 2026: p. 19373341261449883.

35. Marazzi, F., et al., Diagnosis and Treatment of Bone Metastases in Breast Cancer: Radiotherapy, Local Approach and Systemic Therapy in a Guide for Clinicians. Cancers (Basel), 2020. 12(9).

36. Woo, S.M., et al., Cell line IDG-SW3 replicates osteoblast-to-late-osteocyte differentiation in vitro and accelerates bone formation in vivo. J Bone Miner Res, 2011. 26(11): p. 2634–46.

37. Xu, L.H., et al., OCY454 Osteocytes as an in Vitro Cell Model for Bone Remodeling Under Mechanical Loading. J Orthop Res, 2019. 37(8): p. 1681–1689.

38. Aziz, A.H., et al., IDG-SW3 Osteocyte Differentiation and Bone Extracellular Matrix Deposition Are Enhanced in a 3D Matrix Metalloproteinase-Sensitive Hydrogel. ACS Appl Bio Mater, 2020. 3(3): p. 1666–1680.

39. Chen, K., et al., High Mineralization Capacity of IDG-SW3 Cells in 3D Collagen Hydrogel for Bone Healing in Estrogen-Deficient Mice. Front Bioeng Biotechnol, 2020. 8: p. 864.

40. Hoppock, G.A., et al., Connexin 43 and cell culture substrate differentially regulate OCY454 osteocytic differentiation and signaling to primary bone cells. Am J Physiol Cell Physiol, 2023. 325(4): p. C907–C920.

41. Rindt, W.D., et al., A 3D bioreactor model to study osteocyte differentiation and mechanobiology under perfusion and compressive mechanical loading. Acta Biomater, 2024. 184: p. 210–225.

42. Wilmoth, R.L., V.L. Ferguson, and S.J. Bryant, A 3D, Dynamically Loaded Hydrogel Model of the Osteochondral Unit to Study Osteocyte Mechanobiology. Adv Healthc Mater, 2020. 9(22): p. e2001226.

43. Giaquinto, A.N., et al., Breast Cancer Statistics, 2022. CA Cancer J Clin, 2022. 72(6): p. 524–541.

44. Ziegler, P., et al., The impact of physical activity on progression-free and overall survival in metastatic breast cancer based on molecular subtype. BMC Cancer, 2024. 24(1): p. 1284.

45. Maroni, P. and P. Bendinelli, Bone, a Secondary Growth Site of Breast and Prostate Carcinomas: Role of Osteocytes. Cancers (Basel), 2020. 12(7).

46. Florio, M., et al., A bispecific antibody targeting sclerostin and DKK-1 promotes bone mass accrual and fracture repair. Nat Commun, 2016. 7: p. 11505.

47. Pacheco, R. and H. Stock, Effects of radiation on bone. Curr Osteoporos Rep, 2013. 11(4): p. 299–304.

48. Song, X., et al., Vibration-Mediated Recovery of Irradiated Osteocytes and Their Regulatory Role in Breast Cancer Bone Metastasis. Adv Healthc Mater, 2026. 15(4): p. e01689.

