## Supplemental Table 1 for "Novel 3D Mature Osteocyte Perfusion Platform Reveals Subtype-Dependent and Radiation-Sensitive Disruption of Mechanotransduction by Breast Cancer Secretomes"

### SUPPLEMENTAL DATA

**Supplemental Table S1**. Real-time qPCR primer sequences.

| Gene ID | Forward Sequence | Reverse Sequence |
|---|---|---|
| *Sost* (sclerostin) | CCACAAAGACTGAAAGCCGC | TAACAATGCCTCTGGTCGGG |
| *Tnfsf11* (RANKL) | CATTTGCACACCTCACCATCAAT | GTCTGTAGGTACGCTTCCCG |
| *Tnfsf11b* (OPG) | AGAAGCCACGCAAAAGTGTG | TTCACTTTGGTCCCAGGCAA |
| *Hprt* (Hypoxanthine phosphoribosyltransferase) | TCCCAGCGTCGTGATTAGCG | TCGAGCAAGTCTTTCAGTCCTGT |
